# Drug metabolic activity as a selection factor for pluripotent stem cell-derived hepatic progenitor cells

**DOI:** 10.1101/2023.02.21.529337

**Authors:** Saeko Akiyama, Noriaki Saku, Shoko Miyata, Kenta Ite, Hidenori Nonaka, Masashi Toyoda, Akihide Kamiya, Tohru Kiyono, Tohru Kimura, Mureo Kasahara, Akihiro Umezawa

**Affiliations:** Center for Regenerative Medicine, National Center for Child Health and Development Research Institute, Tokyo, 157-8535, Japan; Department of Advanced Pediatric Medicine (National Center for Child Health and Development), Tohoku University School of Medicine, Miyagi, Japan; Research team for Geriatric Medicine (Vascular Medicine), Tokyo Metropolitan Institute of Gerontology, Tokyo, 173-0015, Japan; Department of Molecular Life Sciences, Tokai University School of Medicine, 143 Shimokasuya, Isehara, Kanagawa, 259-1193, Japan; Project for Prevention of HPV-related Cancer, Exploratory Oncology Research and Clinical Trial Center, National Cancer Center, Chiba, 277-8577, Japan; Laboratory of Stem Cell Biology, Department of BioSciences, Kitasato University School of Science, Kanagawa, 252-0373, Japan; Organ Transplantation Center, National Center for Child Health and Development, Tokyo, 157-8535, Japan

**Keywords:** Hepatocyte, Hepatic progenitor cell, Liver regeneration, Cytochrome P450, Pluripotent stem cell

## Abstract

As a metabolic organ, the liver plays a variety of roles, including detoxification. It has been difficult to obtain stable supplies of hepatocytes for transplantation and for accurate hepatotoxicity determination in drug discovery research. Human pluripotent stem cells, capable of unlimited self-renewal, may be a promising source of hepatocytes. In order to develop a stable supply of embryonic stem cell (ESC)-derived hepatocytes, we have purified human ESC-derived hepatic progenitor cells with exposure to cytocidal puromycin by using their ability to metabolize drugs. Hepatic progenitor cells stably proliferated at least 2^20-fold over 120 days, maintaining hepatic progenitor cell-like properties. High drug-metabolizing hepatic progenitor cells can be matured into liver cells by suppressing hepatic proliferative signals. The method we developed enables the isolation and proliferation of functional hepatic progenitors from human ESCs, thereby providing a stable supply of high-quality cell resources at high efficiency. Cells produced by this method may facilitate cell therapy for hepatic diseases and reliable drug discovery research.

## INTRODUCTION

The liver is a major metabolic organ and plays diverse roles in gluconeogenesis, lipid metabolism, and detoxification of ammonia and drugs. Liver transplantation is the only radical treatment for liver disease, but there is a shortage of donors and high costs^1,2^. There are also medical concerns because the procedures are highly invasive for both donors and recipients, and only a limited number of patients can be treated, making the development of cell therapy as an alternative to transplantation desirable^3–5^. To evaluate the pharmacokinetics and toxicity of candidate compounds, studies using experimental animals, primary cultured human hepatocytes, and immortalized hepatocytes have been established^6–9^. After being absorbed, many drugs are metabolized and eliminated by the liver. Metabolites produced in this process can be hepatotoxic and can cause liver damage. Predicting hepatotoxicity specific to humans is essential, and inadequate toxicity prediction models can hinder drug development and clinical trials. Evaluating drug metabolism using human hepatocytes is necessary because the ability to metabolize drugs varies widely among species. However, there is a chronic shortage of human hepatocytes due to difficulties in securing donors and in vitro propagation of hepatocytes.

Against this background, stem cell-derived hepatocytes have recently attracted much attention. Embryonic stem cells (ESCs)^10–12^ and induced pluripotent stem cells (iPSCs) ^13^ can self-renew almost indefinitely while maintaining pluripotency and have been proposed as a stable source of cells such as hepatocytes, which are in high demand in cell therapy, regenerative medicine, and drug discovery research. There are numerous reports of hepatocyte differentiation using human ESCs and human iPSCs^14–17^. In hepatic differentiation from human pluripotent stem cells, it is essential to mimic the process of liver development in vitro. Many differentiation protocols have been adopted to induce hepatocytes via endodermal cells and hepatic progenitor cells (HPCs) from undifferentiated cells^17,18^. The endoderm forms the digestive organs early in development and differentiates into HPCs under signals such as fibroblast growth factor and bone morphogenetic proteins secreted from the adjacent cardiogenic mesoderm. HPCs then proliferate and mature to form the liver under signals such as hepatocyte growth factor. In order to efficiently generate hepatocytes from human pluripotent stem cells, culture systems have been established that utilize HPCs, which have both high proliferative capacity and the ability to differentiate into hepatocytes^19–22^. However, some of these protocols require processes such as cell sorting using specific cell surface markers or gene transfer.

Recently, it has been reported that puromycin treatment is effective in selecting proliferative hepatocytes from the liver of drug-induced liver injury (DILI) patients^23^. Proliferative hepatocytes selected by this method exhibit stable and high proliferative potential in Wnt-containing media and MEF feeders. Furthermore, proliferative hepatocytes possess drug-metabolizing enzyme activities and have a hepatic progenitor cell-like genetic profile, such as co-expression of hepatocyte and biliary epithelial cell markers.

In this study, we aimed to establish an in vitro culture system that can efficiently obtain functional human ESC-derived hepatocytes (ESC-hepatocytes). We successfully isolated HPCs by using puromycin and induced hepatic maturation, which enhances hepatocyte markers and liver function, and suppressed immature hepatocyte markers. This culture system provides an effective and stable supply method for hepatocytes needed for drug discovery research and cell transplantation.

## MATERIALS AND METHODS

### Ethical statement

All experiments handling human cells and tissues were approved by the Institutional Review Board at the National Center for Child Health and Development (IRB No.232). Informed consent was obtained from all participants. When participants were under 18, informed consent was obtained from parents. Human cells in this study were utilized in full compliance with the Ethical Guidelines for Medical and Health Research Involving Human Subjects (Ministry of Health, Labor, and Welfare (MHLW), Japan; Ministry of Education, Culture, Sports, Science and Technology (MEXT), Japan). The derivation and cultivation of ESC lines were performed in full compliance with “the Guidelines for Derivation and Distribution of Human Embryonic Stem Cells (Notification of MEXT, No. 156 of August 21, 2009; Notification of MEXT, No. 86 of May 20, 2010) and “the Guidelines for Utilization of Human Embryonic Stem Cells (Notification of MEXT, No. 157 of August 21, 2009; Notification of MEXT, No. 87 of May 20, 2010)”. Animal experiments were performed in compliance with the basic guidelines for the conduct of animal experiments in implementing agencies under the jurisdiction of the Ministry of Health, Labour and Welfare (Notification of MHLW, No. 0220-1 of February 20, 2015). The protocols of the animal experiments were approved by the Institutional Animal Care and Use Committee of the National Research Institute for Child Health and Development (No. A2003-002-C19-M04). This study was carried out in compliance with the ARRIVE guidelines.

### Preparation of feeder cells

Mouse embryonic fibroblasts (MEFs) were prepared for use as nutritional support (feeder) cells. Heads, limbs, tails, and internal organs were removed from E12.5 ICR mouse fetuses (Japan CLEA, Tokyo, Japan), and the remaining torsos were then minced with a blade and seeded into culture dishes with DMEM supplemented with 10% FBS and 1% penicillin-streptomycin to allow cell growth. After two days of culture, the cells were passaged in a 1:4 ratio. After five days of culture, cells were detached with trypsin, and 1/100 (v/v) of 1M HEPES buffer (15630-106, Invitrogen; Thermo Fisher Scientific, MA, USA) was added to the collected cells. Following irradiation with an X-ray apparatus (dose: 30 Gy, MBR-1520 R-3, Hitachi, Tokyo, Japan), the cells were frozen using a TC protector (TCP-001DS, Pharma Biomedical, Osaka, Japan).

### Human ESC culture

SEES2 ESCs^24^ were cultured on irradiated MEFs (irrMEFs) with medium for human ESCs: Knockout-Dulbecco’s modified Eagle’s medium (KO-DMEM: 10829-018, Life Technologies; Thermo Fisher Scientific, MA, USA) supplemented with 20% Knockout-Serum Replacement (KO-SR: 10828-028, Gibco; Thermo Fisher Scientific, MA, USA), 2 mM Glutamax-I (35050-079, Gibco; Thermo Fisher Scientific, MA, USA), 0.1 mM non-essential amino acids (NEAA) (11140-076, Gibco; Thermo Fisher Scientific, MA, USA), 1% penicillin-streptomycin (Invitrogen) 1 mM sodium pyruvate (11360070, Gibco; Thermo Fisher Scientific, MA, USA), and recombinant human full-length bFGF (PHG0261, Gibco; Thermo Fisher Scientific, MA, USA) at 50 ng/ml^24^.

### Hepatic differentiation of human ESCs

To differentiate ESCs into hepatocytes, embryoid bodies (EBs) were generated by 3D culture to differentiate into endoderm. To generate EBs, ESCs (5 × 10^4 cells/well) were dissociated into single cells with 0.5mM EDTA (15575020, Gibco; Thermo Fisher Scientific, MA, USA) after exposure to ROCK inhibitor, Y-27632 (A11105-01, Fujifilm Wako Pure Chemicals, Osaka, Japan), and cultivated in 96-well plates (174925, Thermo Fisher Scientific, MA, USA) in EB medium [75% KO-DMEM, 20% KO-SR, 2 mM GlutaMAX-I, 0.1 mM NEAA, 1% penicillin-streptomycin] for 4 days. The EBs were transferred to the 24-well plates coated with collagen type I (24 EBs/well) and cultured in XF32 medium [85% Knockout DMEM, 15% Knockout Serum Replacement XF CTS (XF-KSR: 12618013, Gibco; Thermo Fisher Scientific, MA, USA), 2 mM GlutaMAX-I, 0.1 mM NEAA, 1% penicillin-streptomycin, 50 μg/mL L-ascorbic acid 2-phosphate, 10 ng/mL heregulin-1β, 200 ng/mL recombinant human IGF-1 (LONG R3-IGF-1: 85580C, Sigma-Aldrich, MO, USA), and 20 ng/mL human bFGF] for 35±3 days^25^. Cells were detached with Liberase (5339880001, Roche, Basel, Switzerland) and stored frozen at 1×10^7 cells/mL in a Stem Cell Banker (CB047, Nippon Zenyaku Kogyo, Fukushima, Japan) until use.

### Culture and passaging of ESC-hepatocytes

Cryopreserved human ESC-hepatocytes were used. Frozen cells were thawed and seeded onto irrMEFs in 60 mm dishes (3010-060, IWAKI; AGC Techno Glass, Tokyo, Japan) at the seeding density of 5.0 × 10^5 cells/cm^2^ (Passage 1). Then the cells were cultured at 37°C, 5% CO2 with EMUKK-05 medium (Emukk LLC, Japan) containing Wnt3a and R-spondin 1^26^. The medium was changed every three days. After 10-12 days of culture, the cells were trypsinized with 0.25% trypsin-EDTA (23315, IBL, Gunma, Japan) and plated in dishes seeded with irrMEFs in a 1:4 ratio.

### Culture of hepatic progenitor cells from human ESC-derived hepatocytes

For HPCs selection, puromycin (final concentration: 2 μg/mL, 160-23151, Fujifilm Wako Pure Chemicals, Osaka, Japan) was added to 50% confluence of passaged ESC-hepatocytes for three days. After exposure to puromycin, cells were washed with PBS (14190-250, Invitrogen; Thermo Fisher Scientific, MA, USA) and cultured in the EMUKK-05 medium (Emukk LLC, Japan) for at least one day. When the cells reached 90% confluence, the cells were trypsinized with 0.25% trypsin-EDTA (23315, IBL, Gunma, Japan) and plated in dishes seeded with irrMEFs in a 1:4 ratio. At each passaging, the cells were passaged into dishes seeded with irrMEFs, treated with 2 μg/mL puromycin for three days, and cultured without puromycin for at least one day.

### Calculation of population doublings

Cells were harvested at sub-confluency, and the total number of cells in each well was determined using a cell counter. Population doubling (PD) was calculated from the formula PD=log2(A/B), where A is the number of harvested cells and B is the number of plated cells. Population doubling was used as the measure of cell growth.

### Histology and Periodic Acid Schiff (PAS) stain

Samples were suspended in iPGell (PG20-1, GenoStaff, Tokyo, Japan) following the manufacturer’s instructions and fixed in 4□ paraformaldehyde at 4□ overnight. Fixed samples were embedded in a paraffin block to prepare cell sections. For Hematoxylin Eosin (HE) staining, the deparaffinized sections were treated with a hematoxylin solution (Mutoh Chemical, Tokyo, Japan) for 5 min at room temperature and washed with dilute ammonia. After washing with 95% ethanol, dehydration was performed with 150 mL of eosin in 95% ethanol solution and permeabilized in xylene. For PAS staining, the deparaffinized sections were reacted with 0.5% periodate solution (86171, Mutoh Chemical, Tokyo, Japan) for 10 min at room temperature and rinsed with water for 7 min. After reacting with Schiff’s reagent (40922, Mutoh Chemical, Tokyo, Japan) for 5-15 min, the sections were washed with sulfurous acid water. Coloration was achieved by reaction with Meyer hematoxylin solution (30002, Mutoh Chemical, Tokyo, Japan) for 2 min at room temperature and then rinsed with water for 10 min.

### qRT-PCR

Total RNA was prepared using ISOGEN (311-02501, Nippon Gene, Tokyo, Japan) and RNeasy Micro Kit (74004, Qiagen, Hilden, Germany). The RNA was reverse transcribed to cDNA using Superscript □ Reverse Transcriptase (18080-085, Invitrogen; Thermo Fisher Scientific, MA, USA) with ProFlex PCR System (Applied Biosystems, MA, USA). Quantitative RT-PCR was performed on QuantStudio 12K Flex (Applied Biosystems, MA, USA) using a Platinum SYBR Green qPCR SuperMix-UDG (11733046, Invitrogen; Thermo Fisher Scientific, MA, USA). Expression levels were normalized with the reference gene, UBC. The primer sequences are shown in Table 1.

**Table 1.**
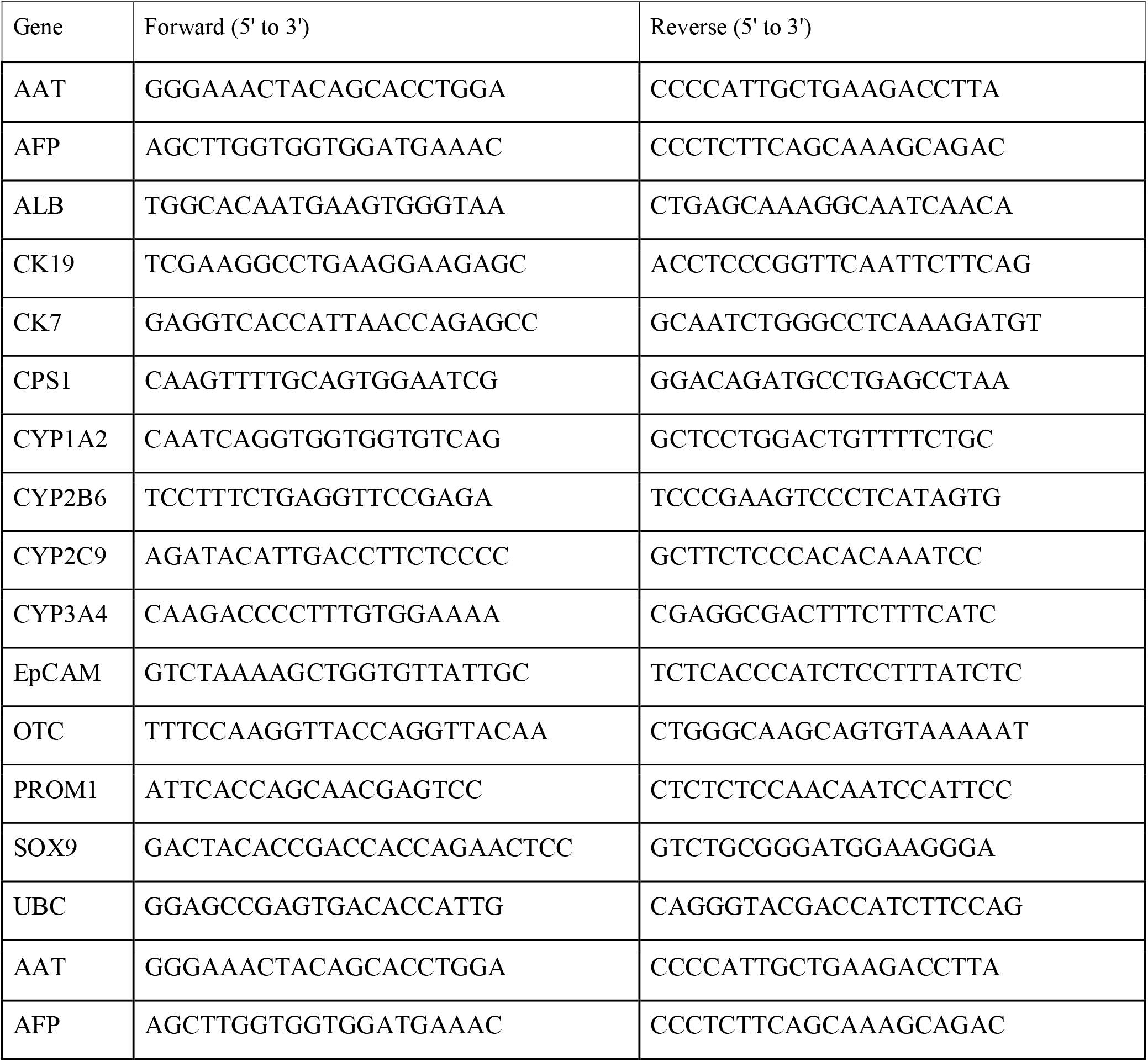
Primers used for qRT-PCR.

### Immunofluorescence staining

Cells were fixed with 4% paraformaldehyde in PBS for 10 min at room temperature. After washing with PBS, cells were permeabilized with 0.1% Triton X-100 in PBS for 10 min, pre-incubated with Protein Block Serum-Free (X0909, Dako, Jena, Germany) for 30 min at room temperature, and then exposed to primary antibodies overnight at 4°C. Cells were washed with PBS and incubated with diluted secondary antibodies for 30 min at room temperature. Nuclei were stained with 4’,6-diamidino-2-phenylindole, and dihydrochloride (DAPI, 40043, Biotium, CA, America). The antibodies (Tables 2 and 3) were diluted according to the tables in PBS containing 1% BSA (126575, Calbiochem).

**Table 2.**
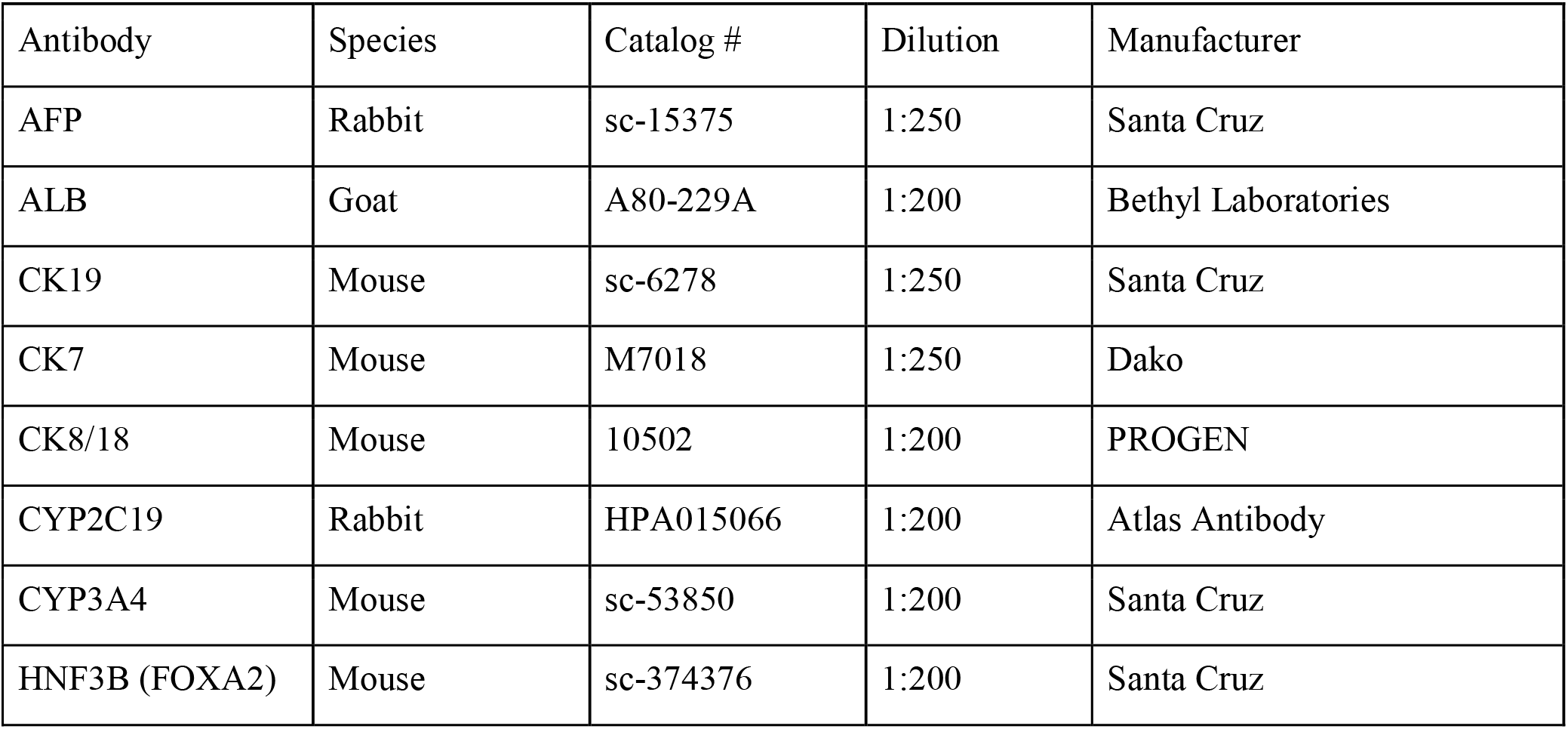
Antibodies used for immunocytochemistry.

**Table 3.** Secondary antibodies used for immunocytochemistry.

| Antibody | Manufacturer | Catalog # |
| --- | --- | --- |
| Alexa Fluor 488 rabbit anti-goat IgG | Invitrogen | A11078 |
| Alexa Fluor 488 goat anti-rabbit IgG | Invitrogen | A11008 |
| Alexa Fluor 488 goat anti-mouse IgG | Invitrogen | A21121 |
| Alexa Fluor 546 goat anti-mouse IgG1 | Invitrogen | A21123 |
| Alexa Fluor 568 goat anti-mouse IgG2a | Invitrogen | A21134 |

### Cytochrome P450 induction

To examine the induction of cytochrome P450 enzymes, cells were cultured in 12-well plates (353043, BD Falcon; Corning, NY, USA) using irrMEFs and EMUKK-05 medium. When the cells reached 80-90% confluence, the following drugs were added to the cells: 20 μM rifampicin (solvent: dimethyl sulfoxide (DMSO), 189-01001, Fujifilm Wako Pure Chemicals, Osaka, Japan) with an induction period of two days, 50 μM omeprazole (solvent: DMSO, 158-03491, Fujifilm Wako Pure Chemicals, Osaka, Japan) with an induction period of one day, and 500 μM phenobarbital (solvent: DMSO, 162-11602, Fujifilm Wako Pure Chemicals, Osaka, Japan) with an induction period of two days.

### Hepatic maturation

For hepatic maturation, ESCs were differentiated into hepatocytes for 35 days in collagen-coated 12-well plates (353043, BD Falcon; Corning, NY, USA). HPCs were cultured in 12-well plates (353043, BD Falcon; Corning, NY, USA) with irrMEFs and EMUKK-05 medium until cells were 80-90% confluent. The following small molecules were then added to each medium; 5 μM DAPT (solvent: DMSO, 049-33583, Fujifilm Wako Pure Chemicals, Osaka, Japan), 10 μM SB431542 (solvent: DMSO, 037-24293, Fujifilm Wako Pure Chemicals, Osaka, Japan), 0.5 μM IWP2 (solvent: DMSO 034-24301, Fujifilm Wako Pure Chemicals, Osaka, Japan), 20 μM forskolin (solvent: DMSO, 063-02193, Fujifilm Wako Pure Chemicals, Osaka, Japan) and 1 μM LDN193189 (solvent: DMSO, 6053-10, Tocris Bioscience Bristol United Kingdom), with an induction period of 10 days.

### Microarray analysis

Total RNA was isolated using miRNeasy mini kit (217004, Qiagen, Hilden, Germany). RNA samples were labeled and hybridized to a SurePrint G3 Human GEO microarray 8 × 60K Ver 3.0 (Agilent, CA, USA), and the raw data were normalized using the 75-percentile shift. Unsupervised clustering was performed with sorted genesets using the R package. Gene set enrichment analysis (GSEA) was performed using gene sets obtained from the Molecular Signatures Database (MSigDB) to examine statistically significant differences in biological status between ESC-hepatocytes and HPCs^27,28^. Gene expression profiles of mature hepatocytes were analyzed using human liver total RNA (636531, Clontech: Takara Bio, Shiga, Japan).

### Statistical analysis

The number of biological and technical replicates is shown in the figure legends. All data are presented as mean ± SD (technical triplicates) or mean ± SE (biological triplicates). In the statistical evaluation of this study, Student’s t-test without correspondence was used to calculate statistical probabilities in the case of two groups, and the Dunnett test was used as a multiple comparison test in the case of three groups. P-values were calculated with a two-tailed t-test.

## RESULTS

### Isolation of hepatic progenitor cells from human ESC-derived hepatocytes

In previous studies, we achieved efficient proliferation by co-culturing irrMEFs with hepatocytes derived from patients with liver disease^23^. When human ESC-hepatocytes were passaged and co-cultured with irrMEFs, the cells formed colonies. Hepatic progenitor cells (HPCs) were selected by adding puromycin to the passaged ESC-hepatocytes (Figures 1A and 1B). Puromycin-selected HPCs formed colonies within a few days after seeding and proliferated until they reached confluency; HPCs proliferated in monolayers and were small in cell size. HPCs showed a high nucleus-cytoplasm ratio, and the nuclei were oval, with no cells having round nuclei characteristic of hepatocytes (Figure 1C). Glycogen accumulation was observed, but to a very low degree (Figure 1C). Immunocytochemical analysis showed that HPCs were positive for the proliferation marker Ki67 and proliferated more than 2^20-fold in 120 days (Figures 1D and 1E).

**Figure 1.**
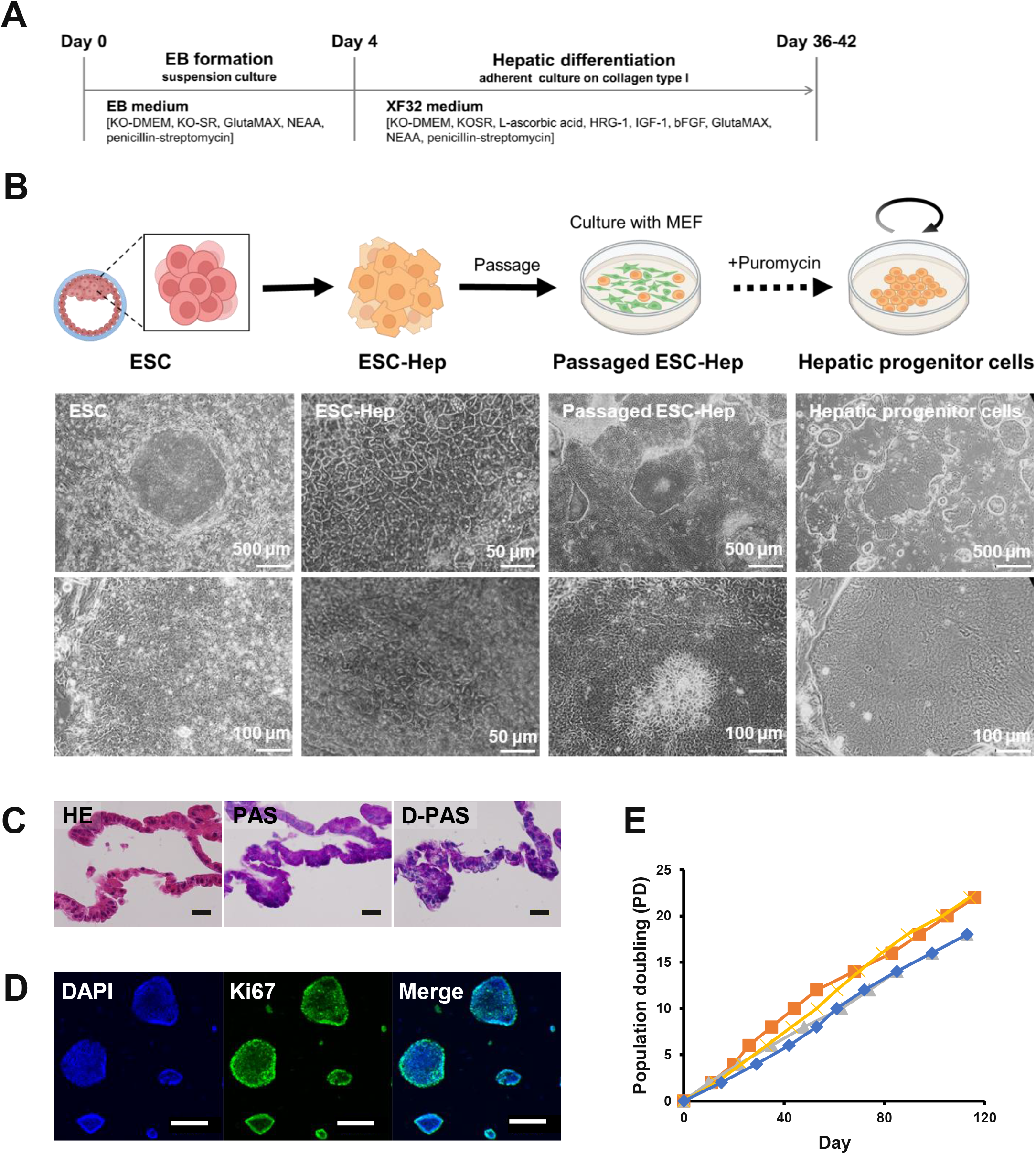
Establishment of hepatic progenitor cells (HPCs) from human ESC-derived hepatocytes. A. Schematic of differentiation protocol from ESCs to hepatocytes B. Scheme of this study and phase-contrast photomicrographs of ESCs, ESC-hepatocytes (ESC-Hep), passaged ESC-hepatocytes (Passaged ESC-Hep), and hepatic progenitor cells. C. Micrographs of HPCs in iPGell. From left to right: HE stain, PAS stain, PAS stain with diastase digestion. D. Immunofluorescence for Ki67 of HPCs. E. Growth curves of HPCs. Cells were passaged in the ratio of 1:4 for each passage. “Population doubling” indicates the cumulative number of divisions of the cell population. Each growth curve in different colors was obtained from independent tetraplicate experiments.

Next, to characterize puromycin-selected HPCs, we analyzed the expression of mature and immature hepatocyte marker genes in ESC-hepatocytes, in passaged ESC-hepatocytes and in puromycin-selected HPCs (Figure 2). The expression of Albumin (ALB), a mature hepatocyte marker, was significantly suppressed in passaged ESC-hepatocytes and in HPCs compared to ESC-hepatocytes, but no difference was observed between passaged ESC-hepatocytes and HPCs. Expression of Hepatocyte nuclear factor 4-alpha (HNF4A), which is essential for hepatic differentiation, the immature hepatocyte markers Alpha-fetoprotein (AFP) and Prominin-1 (PROM1), and the epithelial marker Epithelial cell adhesion molecule (EpCAM) was significantly enhanced in HPCs compared to ESC-hepatocytes and passaged ESC-hepatocytes. The expression of Cytokeratin 7 (CK7), a biliary epithelial cell marker, was also enhanced in HPCs more than in ESC hepatocytes, but its expression level was higher in the passaged ESC-hepatocytes than in HPCs (Figure 2A). The expression of hepatic function markers, the drug-metabolizing enzymes Cytochrome P450 family 1 subfamily A member 2 (CYP1A2), Cytochrome P450 family 2 subfamily B member 6 (CYP2B6), and Cytochrome P450 family 3 subfamily A member 4 (CYP3A4), was higher in HPCs than in passaged ESC-hepatocytes. No significant difference in expression was observed between ESC-hepatocytes, but a marked enhancement of CYP3A4 expression, approximately 500-fold, was observed in HPCs. The urea cycle-related enzyme Carbamoyl phosphate synthetase 1 (CPS1) was more highly expressed in passaged ESC-hepatocytes and HPCs than in ESC-hepatocytes, but there was no difference between passaged ESC-hepatocytes and HPCs. Also, no difference in Ornithine transcarbamylase (OTC) was observed between any of the cells (Figure 2B). Immunocytochemistry showed that the HPCs co-expressed the immature hepatocyte marker AFP, the biliary epithelial cell markers Cytokeratin 19 (CK19) and CK7, and HNF4A, which directs liver characteristics (Figures 2C, 2D, 2E). The hepatic progenitor markers SRY-box9 (SOX9) and Hepatocyte nuclear factor 3-beta (HNF4B) were expressed evenly throughout the colony (Figure 2F). We also confirmed the expression of the hepatic markers ALB and Cytokeratin 8/18 (CK8/18) (Figure 2G) and the drug-metabolizing enzymes CYP3A4 and Cytochrome P450 family 2 subfamily C member 19 (CYP2C19) (Figure 2H). These results indicate that puromycin-selected HPCs have both biliary epithelial cell and hepatocyte characteristics of HPCs and maintain a high and stable proliferative capacity for more than 120 days. Given the markedly enhanced expression of drug-metabolizing enzyme genes in puromycin-selected HPCs, we next evaluated CYP induction in HPCs (Figure 2I). The expression of CYP1A2 was 11.9-fold at 24 h after omeprazole treatment, CYP2B6 was 2.1-fold at 48 h after phenobarbital treatment, and CYP3A4 was 2.1-fold at 48 h after rifampicin treatment, indicating that drug-metabolizing enzymes are significantly increased by the inducers.

**Figure 2.**
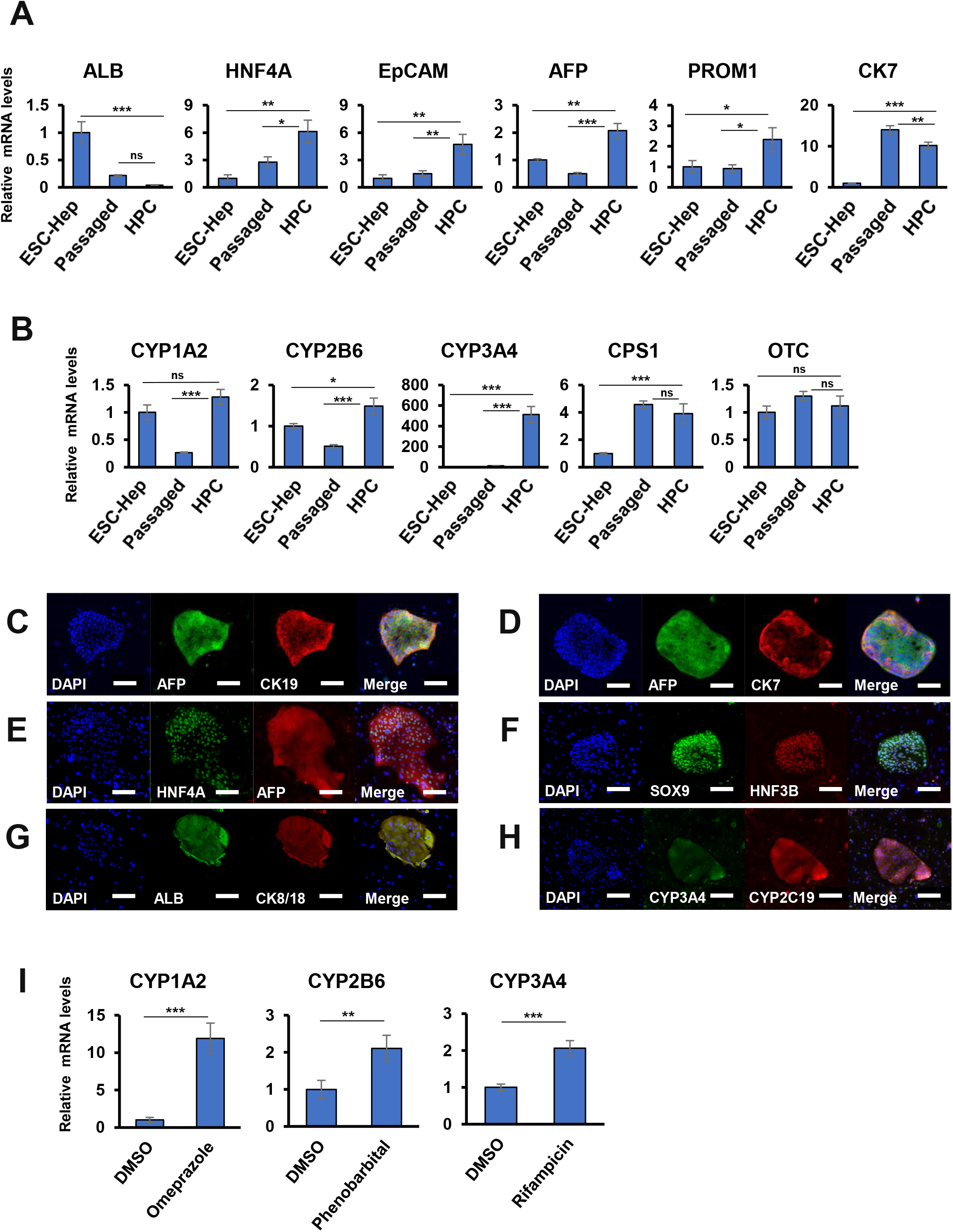
Characteristic analysis of puromycin-selected hepatic progenitor cells (HPCs) A. Gene expression of mature and immature hepatocyte marker genes in passaged ESC-hepatocytes (Passaged) and puromycin-selected HPCs (HPC) analyzed by qRT-PCR. Data were normalized by housekeeping UBC genes and the expression level of each gene in ESC-hepatocytes (ESC-Hep) was set to 1.0. Error bars indicate standard deviation. Statistical analysis was performed by Dunnett’s multiple comparison test. *p < 0.05, **p < 0.01, ***p < 0.001. B. Gene expression of hepatic function marker genes in in passaged ESC-hepatocytes (Passaged) and puromycin-selected HPCs (HPC) analyzed by qRT-PCR. Data were normalized by housekeeping UBC genes, and the expression level of each gene in ESC-Hep was set to 1.0. Error bars indicate standard deviation. Statistical analysis was performed by Dunnett’s multiple comparison test. *p < 0.05, ***p < 0.001. C-H. Immunofluorescence of HPCs for AFP and CK19 (C), AFP and CK7 (D), HNF4A and AFP (E), SOX9 and HNF3B (F), ALB and CK8/18 (G), and CYP3A4 and CYP2C19 (H). I. Induction of drug-metabolizing enzymes in HPCs. Gene expression of CYP1A2, CYP2B6, and CYP3A4 in HPCs after exposure to omeprazole, phenobarbital, and rifampicin was analyzed. The expression level of each gene at no treatment (DMSO) was set at 1.0. Each expression level was calculated from the results of independent (biological) triplicate experiments. Error bars indicate standard error. Student’s T-test was used for statistical analysis of the two groups. **p < 0.01, ***p < 0.001.

### Characterization of puromycin-selected HPCs and human ESC-derived hepatocytes by global gene expression analysis

The gene expression profiles of puromycin-selected HPCs were compared with ESC-hepatocytes and fresh mature hepatocytes isolated from livers (‘Liver’) (Figure 3). Principal component analysis revealed that HPCs were classified into an independent group different from ESC-hepatocytes and ‘Liver’, regardless of the number of passages (Figure 3A). Fetal hepatobiliary progenitor cell-related genes were strongly expressed in HPCs compared to ESC-hepatocytes (Figure 3B); the expression pattern of HPCs was maintained over passaging and was different from that of ESC-hepatocytes. These results suggest that puromycin-selected HPCs have more stable characteristics similar to hepatic progenitor cells in human fetal liver than ESC-hepatocytes. To further characterize puromycin-selected HPCs and ESC-hepatocytes in liver function, we compared gene expression between HPCs and ESC-hepatocytes and identified pathways whose expression was increased in HPCs by Gene Set Enrichment Analysis. The results showed that the gene sets related to drug metabolism were highly enriched (FDR < 0.25, p-value < 0.05) in HPCs compared to ESC-hepatocytes (Figure 3C, Table 4). The heat map shows that HPCs had a higher expression of a set of genes related to drug metabolism than ESC-hepatocytes, regardless of passages (Figure 3D). Steroid hormone biosynthesis was significantly enriched in HPCs compared to ESC-hepatocytes. No significant differences were observed for other hepatic functions, although there was a trend toward enrichment HPCs (Figure 3E, Table 4).

**Figure 3.**
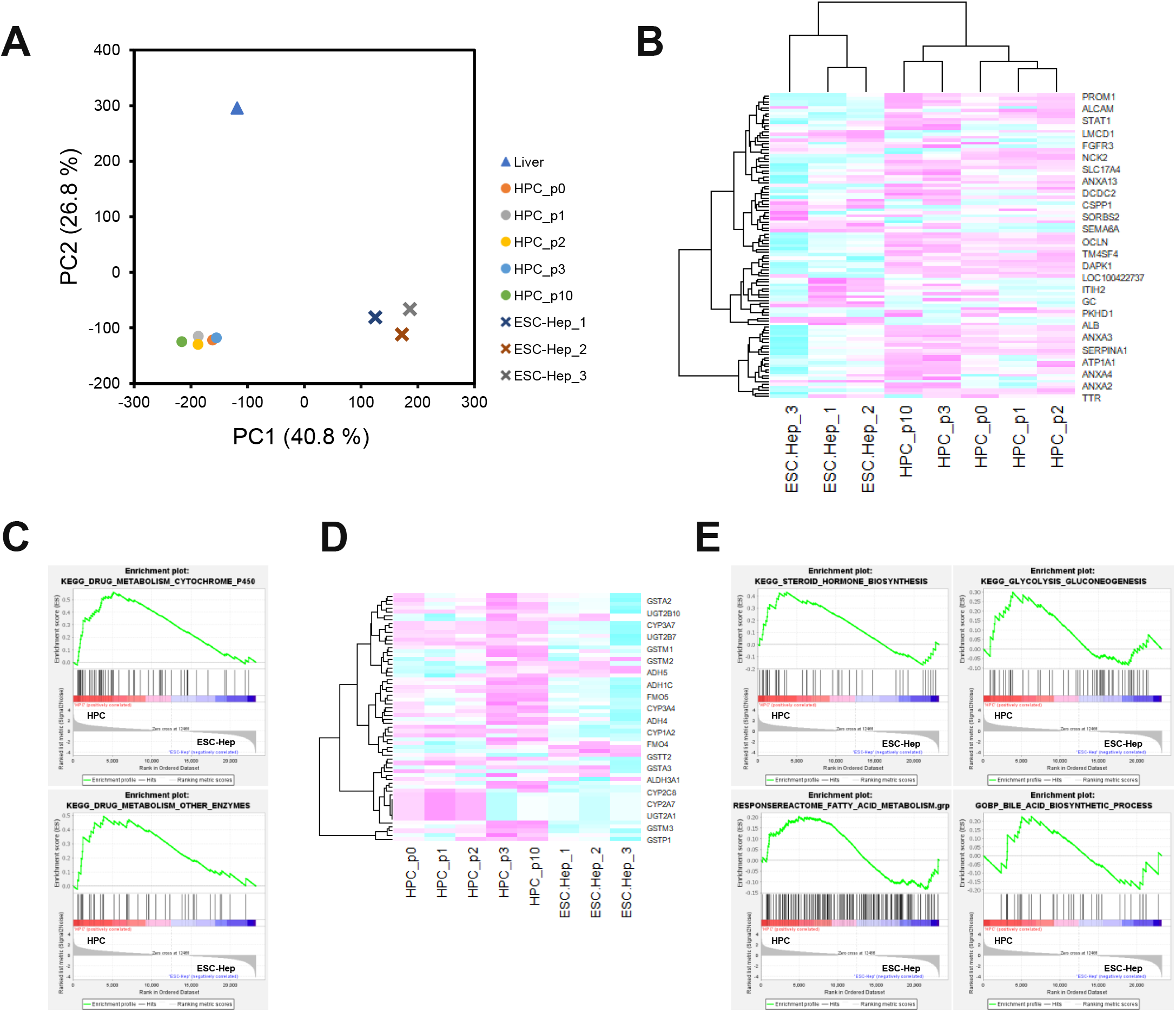
Global gene expression analysis reveals profiles of ESC-hepatocytes (ESC-Hep) and hepatic progenitor cells (HPCs) A. Principal-component analysis (PCA) for HPCs at passage 0, 1, 2, 3 and 10 (HPC_p0, HPC_p1, HPC_p2, HPC_p3, HPC_p10), ESC-hepatocytes (ESC-Hep_1, ESC-Hep_2, ESC-Hep_3), and mature hepatocytes isolated from livers (Liver) by global gene expression. B. Heatmap of the fetal hepatobiliary progenitor up-regulated genes (gene list: https://www.nature.com/articles/s41467-019-11266-x) in HPCs and ESC-hepatocytes. The color bars show the signal strength scaled by the z-score. C. Gene Set Enrichment Analysis (GSEA) of drug metabolism-related genes in HPCs and ESC-Hep. The gene set collection used for the analysis was obtained from KEGG. The cutoff for significant gene sets was FDR < 0.25. Details are shown in Table 4. D. Heatmap of the drug metabolism-associated genes in HPCs and ESC-hepatocytes. Color bars indicate signal intensity scaled by z-score. E. Gene Set Enrichment Analysis (GSEA) for liver gene sets upregulated in HPCs compared to ESC-Heps. Gene set collections used for analysis were obtained from KEGG, Reactome, or GO. The cutoff for significant gene sets was FDR < 0.25. Details are shown in Table 4.

**Table 4.**
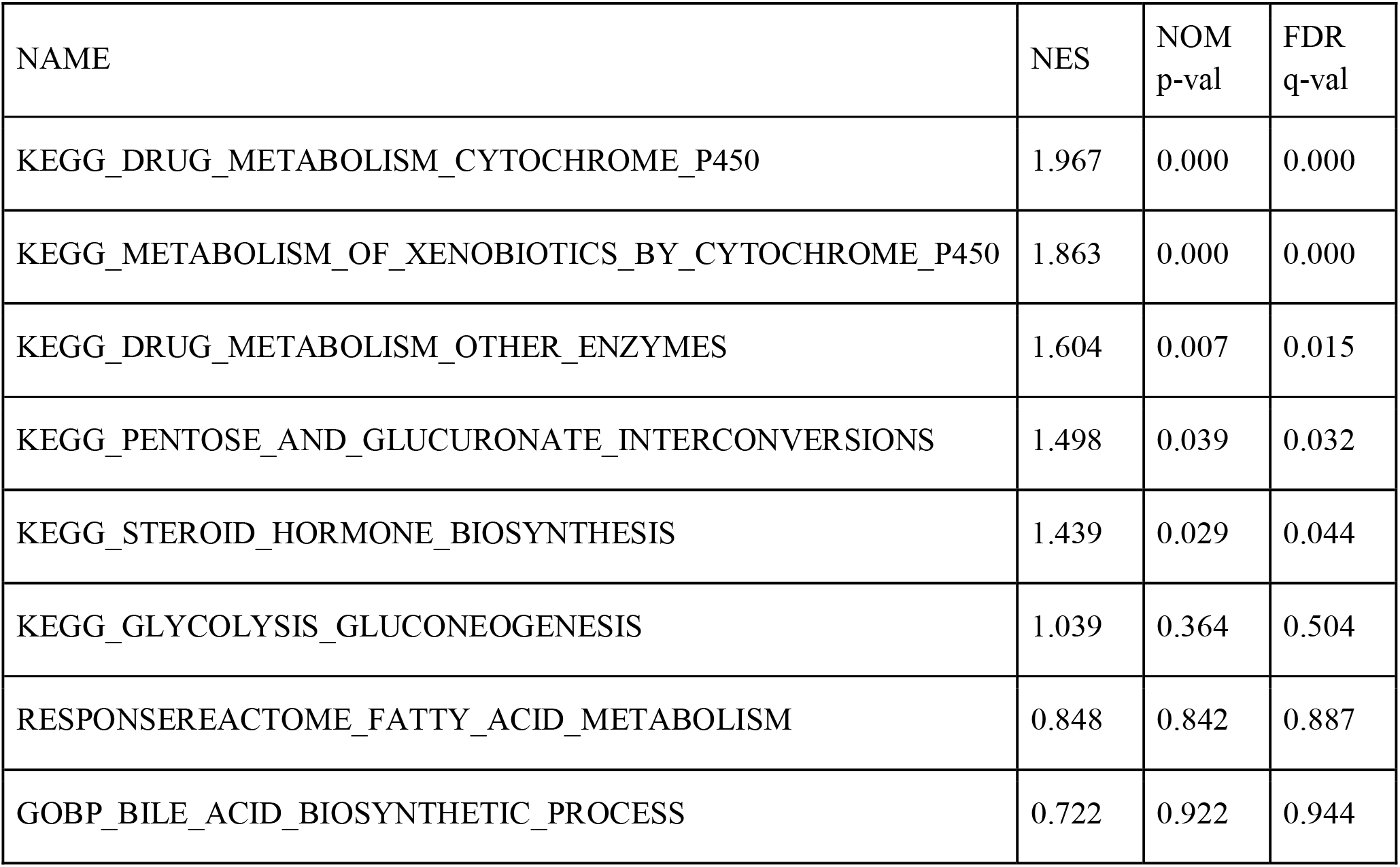
GSEA analysis results that are upregulated in HPCs compared to ESC-hepatocytes.

### Maturation of ESC-hepatocytes and HPCs by 5 compounds

We investigated whether puromycin-selected HPCs acquired the properties of hepatocytes. Since global gene expression analysis showed that ESC-hepatocytes and HPCs have different properties from mature hepatocytes, we first examined the substances effective for further hepatic maturation of ESC-hepatocyte: Wnt inhibitor, Notch inhibitor, forskolin, BMP inhibitor, and TGFb inhibitor. Five types of compounds (5C) were added to ESC-hepatocytes 35 days after the end of differentiation. Solvent DMSO treatment had no significant effect on gene expression in ESC-hepatocytes, and the mature hepatocyte marker ALB and the drug-metabolizing enzymes CYP1A2 and CYP3A4 were significantly enhanced only when 5C was added. No significant differences were observed for the immature hepatocyte marker AFP, the biliary epithelial cell marker CK7, and urea cycle-related enzymes (Figure 4A). These results indicate that the five compounds are effective in the hepatic maturation of ESC-hepatocytes.

**Figure 4.**
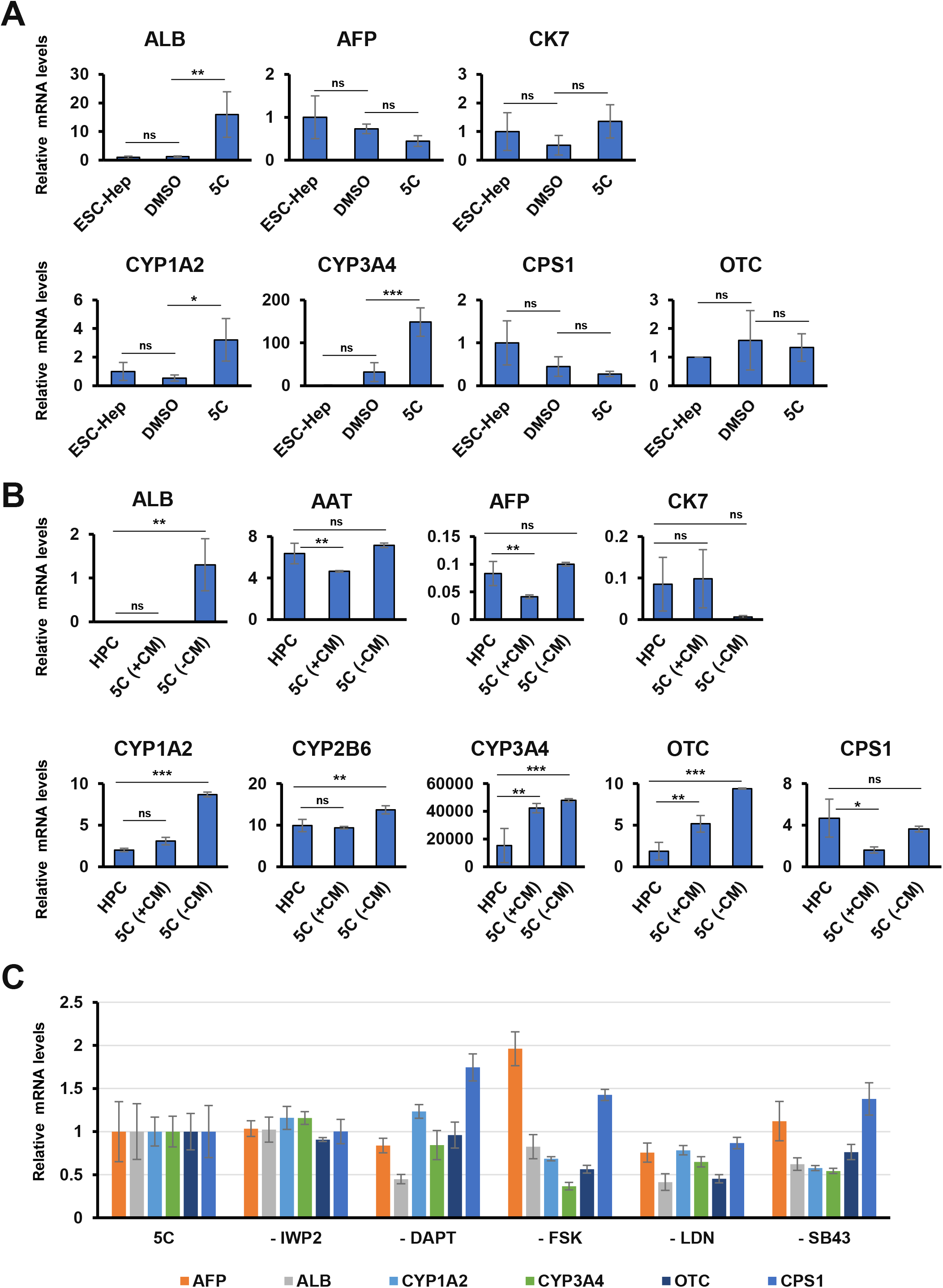
Hepatic progenitor cells (HPCs) are matured by the five compounds (5C) A. ESC-hepatocytes (ESC-Hep) are matured by the five compounds (5C). Gene expression of mature and immature hepatocyte markers in the non-added group (solvent: DMSO) and 5C added group (5C) were analyzed by qRT-PCR. The expression level of each gene in ESC-Hep at day 35, immediately after the end of differentiation induction, was set as 1.0. Each expression level was calculated from the results of independent (biological) duplicate experiments. Error bars indicate standard error. Statistical analysis was performed by Dunnett’s multiple comparison test. *p < 0.05, **p < 0.01, ***p < 0.001. B. HPCs are matured by 5C. Gene expression of mature and immature hepatocyte markers was analyzed by qRT-PCR in control HPCs, in 5C-treated groups with Wnt (5C (+Wnt)), and in 5C-treated groups without Wnt (5C(-Wnt)). The expression level of each gene in the HPCs was set at 1.0. Each expression level was calculated from the results of independent (biological) triplicate experiments. Error bars indicate standard error. Statistical analysis was performed by Dunnett’s multiple comparison test. *p < 0.05, **p < 0.01, ***p < 0.001. C. Effect of the removal of individual compounds from 5C for hepatic maturation in HPCs. Gene expression was analyzed by qRT-PCR in the group with five compounds (5C) and in the group with one compound removed (-IWP2, -DAPT, -FSK, -LDN, -SB43). The expression level of each gene in the 5C-treated group was set at 1.0. Each expression level was calculated from the results of independent (biological) triplicate experiments. Error bars indicate standard errors.

We next investigated whether 5C induced hepatic maturation of puromycin-selected HPCs as well as ESC-hepatocytes; since the medium used for HPCs culture includes Wnt, we decided to investigate the effect of Wnt on HPCs maturation with 5C (Figure 4B). 5C-treated HPCs in a Wnt-free medium significantly enhanced expression of the mature hepatocyte marker ALB, the drug-metabolizing enzymes CYP1A2, CYP2B6, and CYP3A4, and the urea cycle-related enzyme OTC compared to untreated HPCs. No changes were observed in the expression of Alpha-1 antitrypsin (AAT), a mature hepatocyte marker, and AFP, an immature hepatocyte marker. The expression of CK7, a biliary epithelial cell marker, tended to be suppressed. However, compared to ESC-hepatocytes, 5C treatment significantly enhanced the expression of mature hepatocyte markers and liver function markers by more than 10-fold and significantly suppressed the expression of immature hepatocyte markers and biliary epithelial cell markers by less than 1/10. 5C effects were observed only in the absence of Wnt and were suppressed in the presence of Wnt (Figure 4B).

Finally, we investigated which of these five compounds was more effective in the hepatic by comparing the expression of HPCs treated with all five compounds to HPCs with one of the compounds removed. We found that HPCs from which the Notch inhibitor (DAPT), forskolin (FSK), BMP inhibitor (LDN193189), and TGFb inhibitor (SB431542) were removed had lower expression of mature hepatocyte markers and liver function markers than the group treated with all 5C (Figure 4C). However, HPCs removed from the Wnt inhibitor IWP2 did not show suppressed expression. These results indicate that puromycin-selected HLCs are capable of restoring hepatocytic properties with Notch inhibitor, BMP inhibitor, TGFb inhibitor, and forskolin.

## DISCUSSION

Human hepatocytes are in increasing demand for applications, including hepatic pathophysiology, drug discovery, cell therapy, and regenerative medicine^29–32^. However, donor hepatocytes are difficult to obtain and have low proliferative potential, making chronic shortages of hepatocytes. Therefore, there is a need for a system to secure and stably acquire hepatocytes using pluripotent stem cells such as ESCs and iPSCs, which have unlimited proliferative capacity while maintaining pluripotency.

In this study, by using puromycin and establishing a stable proliferative culture system, we succeeded in increasing the number of hepatocytic progenitors by at least 2^20-fold in 120 days. Puromycin equalized hepatocytes with diverse functions such as drug metabolism after the in vitro propagation. The stable proliferative capacity of primary hepatocytes and pluripotent stem cell-derived hepatocytes requires WNT, R-sponsin1, and feeder cells^23,33–36^. Resistance to puromycin may be due to the high metabolic activities of CYP3A4. This metabolism of puromycin by CYP3A4 can indeed be predicted by the computer algorithm “admetSAR” and previous reports^23,37,38^.

The HPCs in this study are almost equivalent to human hepatocytic progenitors from viewpoints of proliferative capability and hepatic phenotypes^14–17^.

Five compounds, i.e. Wnt inhibitor, BMP inhibitor, TGFb inhibitor, Notch inhibitor, and forskolin are effective for in vitro maintenance of human hepatocytes^39^. In addition to hepatic maintenance, Wnt, BMP/TGFb, Notch, and cAMP signalings are involved in the maturation of ESC-derived hepatocytic progenitors as shown in this study. Forskolin promotes hepatic maturation by increasing intracellular cAMP levels^40^. Wnt, BMP/TGFb, and Notch signaling are involved in liver regeneration and hepatocyte proliferation^41–45^. In particular, Wnt plays a vital role in hepatocytic proliferation/division, and Wnt is directly involved in hepatocyte proliferation in vivo and in vitro^34,35,46,47^. There are four types of hepatocytes in the regenerating liver: (1) quiescent, (2) transitional, (3) proliferative, and (4) hypermetabolic^48^. Cells in the hypermetabolic state highly express biosynthesis- and metabolism-related genes. In contrast, cells in the proliferative state express genes related to cell division and cell proliferation. Indeed, hepatocyte function genes are mainly expressed in non-proliferating cells, while these are at low levels in replicating cells^49^. These molecular mechanisms in hepatic regeneration and function may lead to the hypothesis that HPCs in this study can be induced in terminally matured hepatocytes with functionality and engraftability by decreasing proliferation signalings.

Since pluripotent stem cell-derived cells are contaminated with non-target cells that follow similar differentiation pathways, there is a need to develop protocols that can give pure, high-yield differentiated target cells. We succeeded in obtaining ESC-derived hepatocytes that exhibit enhanced expression of hepatic markers, suppression of immature liver markers, and suppression of biliary epithelial cell epithelial markers. Improvement of liver-related function is achieved by the enrichment using liver-specific drug metabolism and hepatic maturation by a combination of effective low-molecules. An efficient and selective hepatic progenitor cell proliferation system and a stable supply of hepatocytes through induction of hepatic maturation would benefit the high need for pluripotent stem cell-derived hepatocytes and the establishment of an alternative platform to animal experiments in drug discovery and the widespread use of cell therapy.

Additionally, iPSCs in combination with genome editing technology will enable research and treatment of liver genetic diseases.

## Ethics approval and consent to participate

Human cells in this study were performed in full compliance with the Ethical Guidelines for Clinical Studies (Ministry of Health, Labor, and Welfare, Japan). The cells were banked after approval of the Institutional Review Board at the National Institute of Biomedical Innovation (May 9, 2006). Animal experiments were performed according to protocols approved by the Institutional Animal Care and Use Committee of the National Research Institute for Child Health and Development.

## Consent for publication

Not applicable.

## Availability of data and materials

The datasets used and analyzed during the current study are available from the corresponding author upon reasonable request.

## Author Contribution

AU and SA designed the experiments. SA, NS, SM and KI performed the experiments. SA and AU analyzed data. HN, MT, AK, and TKiy contributed to the reagents and tissues. SA, HN, MT, AK, TKiy, TKim, MK, and AU discussed the data and manuscript. SA and AU wrote this manuscript.

## Acknowledgments

We would like to express our sincere thanks to K. Miyado and H. Akutsu for the fruitful discussion, M. Ichinose and the HAES team for providing expert technical assistance, and C. Ketcham for English editing and proofreading, and to E. Suzuki and K. Saito for secretarial work. □

## REFERENCES

1. Kasahara M, Katono M, Schlegel A, et al. Waiting list mortality for pediatric deceased donor liver transplantation in a Japanese living-donor-dominant program. Pediatr Transplant. 2019;23(8):e13578.

2. Kim WR, Lake JR, Smith JM, et al. OPTN/SRTR 2017 Annual Data Report: Liver. Am J Transplant. 2019;19 Suppl 2:184–283.

3. Nagasue N, Kohno H, Matsuo S, et al. Segmental (partial) liver transplantation from a living donor. Transplant Proc. 1992;24(5):1958–1959.

4. Cheah YL, Simpson MA, Pomposelli JJ, Pomfret EA. Incidence of death and potentially life-threatening nearmiss events in living donor hepatic lobectomy: a world-wide survey. Liver Transpl. 2013;19(5):499–506.

5. Dhawan A, Puppi J, Hughes RD, Mitry RR. Human hepatocyte transplantation: current experience and future challenges. Nat Rev Gastroenterol Hepatol. 2010;7(5):288–298.

6. Strom SC, Davila J, Grompe M. Chimeric mice with humanized liver: tools for the study of drug metabolism, excretion, and toxicity. Methods Mol Biol. 2010;640:491–509.

7. Zhang D, Luo G, Ding X, Lu C. Preclinical experimental models of drug metabolism and disposition in drug discovery and development. Yao Xue Xue Bao. 2012;2(6):549–561.

8. Yokoyama Y, Sasaki Y, Terasaki N, et al. Comparison of Drug Metabolism and Its Related Hepatotoxic Effects in HepaRG, Cryopreserved Human Hepatocytes, and HepG2 Cell Cultures. Biol Pharm Bull. 2018;41(5):722–732.

9. Peltz G. Can “humanized” mice improve drug development in the 21st century? Trends Pharmacol Sci. 2013;34(5):255–260.

10. Evans MJ, Kaufman MH. Establishment in culture of pluripotential cells from mouse embryos. Nature. 1981;292(5819):154–156.

11. Martin GR. Isolation of a pluripotent cell line from early mouse embryos cultured in medium conditioned by teratocarcinoma stem cells. Proceedings of the National Academy of Sciences. 1981;78(12):7634–7638.

12. Thomson JA, Itskovitz-Eldor J, Shapiro SS, et al. Embryonic stem cell lines derived from human blastocysts. Science. 1998;282(5391):1145–1147.

13. Takahashi K, Yamanaka S. Induction of Pluripotent Stem Cells from Mouse Embryonic and Adult Fibroblast Cultures by Defined Factors. Cell. 2006;126(4):663–676.

14. Corbett JL, Duncan SA. iPSC-Derived Hepatocytes as a Platform for Disease Modeling and Drug Discovery. Front Med. 2019;6:265.

15. Rambhatla L, Chiu CP, Kundu P, Peng Y, Carpenter MK. Generation of hepatocyte-like cells from human embryonic stem cells. Cell Transplant. 2003;12(1):1–11.

16. Schwartz RE, Trehan K, Andrus L, et al. Modeling hepatitis C virus infection using human induced pluripotent stem cells. Proc Natl Acad Sci U S A. 2012;109(7):2544–2548.

17. Ang LT, Tan AKY, Autio MI, et al. A Roadmap for Human Liver Differentiation from Pluripotent Stem Cells. Cell Rep. 2018;22(8):2190–2205.

18. Cai J, Zhao Y, Liu Y, et al. Directed differentiation of human embryonic stem cells into functional hepatic cells. Hepatology. 2007;45(5):1229–1239.

19. Takayama K, Nagamoto Y, Mimura N, et al. Long-term self-renewal of human ES/iPS-derived hepatoblast-like cells on human laminin 111-coated dishes. Stem Cell Reports. 2013;1(4):322–335.

20. Yanagida A, Ito K, Chikada H, Nakauchi H, Kamiya A. An in vitro expansion system for generation of human iPS cell-derived hepatic progenitor-like cells exhibiting a bipotent differentiation potential. PLoS One. 2013;8(7):e67541.

21. Zhao D, Chen S, Cai J, et al. Derivation and characterization of hepatic progenitor cells from human embryonic stem cells. PLoS One. 2009;4(7):e6468.

22. Zhang M, Sun P, Wang Y, et al. Generation of Self-Renewing Hepatoblasts From Human Embryonic Stem Cells by Chemical Approaches. Stem Cells Transl Med. 2015;4(11):1275–1282.

23. Akiyama S, Saku N, Miyata S, et al. Drug metabolic activity is a critical cell-intrinsic determinant for selection of hepatocytes during long-term culture. Stem Cell Res Ther. 2022;13(1):104.

24. Akutsu H, Nasu M, Morinaga S, et al. In vivo maturation of human embryonic stem cell-derived teratoma over time. Regen Ther. 2016;5:31–39.

25. Akutsu H, Machida M, Kanzaki S, et al. Xenogeneic-free defined conditions for derivation and expansion of human embryonic stem cells with mesenchymal stem cells. Regen Ther. 2015;1:18–29.

26. Yachida S, Wood LD, Suzuki M, et al. Genomic Sequencing Identifies ELF3 as a Driver of Ampullary Carcinoma. Cancer Cell. 2016;29(2):229–240.

27. Subramanian A, Tamayo P, Mootha VK, et al. Gene set enrichment analysis: a knowledge-based approach for interpreting genome-wide expression profiles. Proc Natl Acad Sci U S A. 2005;102(43):15545–15550.

28. Mootha VK, Lindgren CM, Eriksson KF, et al. PGC-1α-responsive genes involved in oxidative phosphorylation are coordinately downregulated in human diabetes. Nat Genet. 2003;34(3):267–273.

29. Enosawa S, Horikawa R, Yamamoto A, et al. Hepatocyte transplantation using a living donor reduced graft in a baby with ornithine transcarbamylase deficiency: a novel source of hepatocytes. Liver Transpl. 2014;20(3):391–393.

30. Strom SC, Chowdhury JR, Fox IJ. Hepatocyte transplantation for the treatment of human disease. Semin Liver Dis. 1999;19(1):39–48.

31. Iansante V, Mitry RR, Filippi C, Fitzpatrick E, Dhawan A. Human hepatocyte transplantation for liver disease: current status and future perspectives. Pediatr Res. 2018;83(1-2):232–240.

32. Guo L, Dial S, Shi L, et al. Similarities and Differences in the Expression of Drug-Metabolizing Enzymes between Human Hepatic Cell Lines and Primary Human Hepatocytes. Drug Metabolism and Disposition. 2011;39(3):528–538.

33. Tsuneishi R, Saku N, Miyata S, et al. Ammonia-based enrichment and long-term propagation of zone I hepatocyte-like cells. Sci Rep. 2021;11(1):11381.

34. Zhang K, Zhang L, Liu W, et al. In Vitro Expansion of Primary Human Hepatocytes with Efficient Liver Repopulation Capacity. Cell Stem Cell. 2018;23(6):806–819.e4.

35. Katsuda T, Matsuzaki J, Yamaguchi T, et al. Generation of human hepatic progenitor cells with regenerative and metabolic capacities from primary hepatocytes. Elife. 2019;8.

36. Okada K, Kamiya A, Ito K, et al. Prospective Isolation and Characterization of Bipotent Progenitor Cells in Early Mouse Liver Development. Stem Cells and Development. 2012;21(7):1124–1133.

37. Cheng F, Li W, Zhou Y, et al. admetSAR: A Comprehensive Source and Free Tool for Assessment of Chemical ADMET Properties. Journal of Chemical Information and Modeling. 2012;52(11):3099–3105.

38. Miyata S, Saku N, Akiyama S, et al. Puromycin-based purification of cells with high expression of the cytochrome P450 CYP3A4 gene from a patient with drug-induced liver injury (DILI). Stem Cell Res Ther. 2022;13(1):6.

39. Xiang C, Du Y, Meng G, et al. Long-term functional maintenance of primary human hepatocytes in vitro. Science. 2019;364(6438):399–402.

40. Ogawa S, Surapisitchat J, Virtanen C, et al. Three-dimensional culture and cAMP signaling promote the maturation of human pluripotent stem cell-derived hepatocytes. Development. 2013;140(15):3285–3296.

41. Herrera B, Addante A, Sánchez A. BMP Signalling at the Crossroad of Liver Fibrosis and Regeneration. Int J Mol Sci. 2017;19(1).

42. Schaub JR, Huppert KA, Kurial SNT, et al. De novo formation of the biliary system by TGFβ-mediated hepatocyte transdifferentiation. Nature. 2018;557(7704):247–251.

43. Jin Y, Anbarchian T, Wu P, et al. Wnt signaling regulates hepatocyte cell division by a transcriptional repressor cascade. Proc Natl Acad Sci U S A. 2022;119(30):e2203849119.

44. Yanger K, Zong Y, Maggs LR, et al. Robust cellular reprogramming occurs spontaneously during liver regeneration. Genes Dev. 2013;27(7):719–724.

45. Bogaerts E, Heindryckx F, Vandewynckel YP, Van Grunsven LA, Van Vlierberghe H. The roles of transforming growth factor-β, Wnt, Notch and hypoxia on liver progenitor cells in primary liver tumours (Review). Int J Oncol. 2014;44(4):1015–1022.

46. Liu Z, Kuna VK, Xu B, Sumitran-Holgersson S. Wnt ligands 3a and 5a regulate proliferation and migration in human fetal liver progenitor cells. Transl Gastroenterol Hepatol. 2021;6:56.

47. Monga SP, Pediaditakis P, Mule K, Stolz DB, Michalopoulos GK. Changes in WNT/beta-catenin pathway during regulated growth in rat liver regeneration. Hepatology. 2001;33(5):1098–1109.

48. Chembazhi UV, Bangru S, Hernaez M, Kalsotra A. Cellular plasticity balances the metabolic and proliferation dynamics of a regenerating liver. Genome Res. 2021;31(4):576–591.

49. Walesky CM, Kolb KE, Winston CL, et al. Functional compensation precedes recovery of tissue mass following acute liver injury. Nat Commun. 2020;11(1):5785.

